# *Borrelia* infection in rodent host has dramatic effects on the microbiome of ticks

**DOI:** 10.1101/2021.03.15.435198

**Authors:** Phineas T. Hamilton, Elodie Maluenda, Anouk Sarr, Alessandro Belli, Georgia Hurry, Olivier Duron, Olivier Plantard, Maarten J. Voordouw

## Abstract

**Background:** Vector-borne diseases remain major causes of human morbidity and mortality. It is increasingly recognized that the community of microbes inhabiting arthropods can strongly affect their vector competence, but the role of the tick microbiome in *Borrelia* transmission – the cause of Lyme disease – remains unclear.

**Results:** Here, we use a large-scale experiment to clarify the reciprocal interactions between *Borrelia afzelii* and the microbiome of *Ixodes ricinus*, its primary vector. In contrast to other reports, we find that depletion of the bacterial microbiome in larval ticks has no effect on their subsequent acquisition of *B. afzelii* during blood feeding on infected mice. Rather, exposure to *B. afzelii*-infected hosts drives pervasive changes to the tick microbiome, decreasing overall bacterial abundance, shifting bacterial community composition, and increasing bacterial diversity. These effects appear to be independent of the acquisition of *B. afzelii* by ticks, suggesting they are mediated by physiological or immunological aspects of *B. afzelii* infection in the rodent host.

**Conclusions:** Manipulation of the microbiome of *I. ricinus* larvae had no effect on their ability to acquire *B. afzelii*. In contrast, *B. afzelii* infection in the mouse had dramatic effects on the composition of the gut microbiome in *I. ricinus* nymphs. Our study demonstrates that vector-borne infections in the vertebrate host shape the microbiome of the arthropod vector.

## Background

Infectious diseases vectored by arthropods impose an enormous burden on human health [1-3]. For successful transmission, however, vector-borne pathogens must contend with both the immunological defenses of the arthropod vector, and the community of other microbiota that inhabit it [4-7], and it is now clear that endogenous microbes can shape the competence of diverse arthropod vectors in acquiring and transmitting pathogens [8-12]. Experimental perturbations of the microbiome (‘dysbiosis’) via the use of antibiotics or other methods have, in various contexts, been shown to either increase or decrease the susceptibility of arthropods to colonization by vector-borne pathogens [8, 10, 12, 13]. In particular, the introduction of the intracellular bacterium *Wolbachia* into mosquito vectors [14] can dramatically reduce mosquito competence to vector arboviruses, malaria parasites, and filarial nematodes (Kambris et al. 2009, Moreira et al. 2009, Bian et al. 2010). These effects appear to be mediated through the innate immune system of the arthropod host, rendering the mosquito less susceptible to infection by diverse pathogens [15-17]. Similarly, *Enterobacter* bacteria in the midgut of Anopheline mosquitoes produce reactive oxygen species that can kill malaria parasites [9]. Such direct and indirect antagonistic interactions have formed the basis of increasingly sophisticated biological control strategies that include the ongoing use of *Wolbachia* to control Dengue virus transmission by *Aedes* mosquitoes [3, 14, 18, 19].

Hard ticks remain among the most important vectors of infectious disease in the northern hemisphere, transmitting numerous pathogens that include the causative agents of Lyme borreliosis (‘Lyme disease’), anaplasmosis, babesiosis, and tick-borne encephalitis [20-23]. In particular, the increasing incidence of Lyme and other tick-borne diseases in parts of Europe and North America has underscored the public health risks associated with hard ticks [24-28]. Recent work on *Ixodes scapularis* has suggested that perturbations to the tick’s microbiome can influence tick susceptibility to *B. burgdorferi* sensu stricto (ss) and *Anaplasma phagocytophilum* by affecting the integrity of the tick midgut [8, 12], although the generality and importance of these effects in the natural transmission of these pathogens are unclear. It is likewise unclear if and when specific members of the tick microbiota can interfere with *Borrelia* colonization of the tick, or if *Borrelia* impacts the tick microbiota in ways that might affect the dynamics of other tick-borne diseases.

Here, we use a large-scale experiment with tick offspring derived from wild-collected gravid tick mothers to investigate reciprocal interactions between *Borrelia afzelii*, an endemic cause of Lyme disease in Europe [29], and the microbiome of *Ixodes ricinus*, its primary vector. We found that disrupting the microbiome of larval *I. ricinus* ticks by bleaching egg casings has profound yet transient effects on the tick microbiota, but no effect on subsequent colonization by *B. afzelii* when ticks feed on *B. afzelii-*infected hosts. In contrast, feeding on *B. afzelii*-infected mice has pervasive effects on the composition and diversity of the tick microbiome, which are largely independent of prior microbiome disruption. This suggests that *B. afzelii* infection dynamics within rodent hosts have strong potential to sculpt the tick microbiota, with unclear consequences for the transmission of other tick-borne pathogens. Negative interactions between *B. afzelii* and other members of the tick microbiome might provide insights into the biological control of tick-borne diseases.

## Methods

### *Borrelia*, ticks, and mice

We used *Borrelia afzellii* isolate NE4049, which was obtained from an *I. ricinus* nymph at a field site near Neuchâtel, Switzerland. This strain has multi-locus sequence type ST679, *ospC* major group allele A10, and strain ID number 1887 in the *Borrelia* MLST database. We used isolate NE4049 because we have previously shown that it is highly infectious for both mice and ticks [30-32]. Pathogen-free, female *Mus musculus* BALB/c ByJ mice were used as the vertebrate reservoir host. To infect mice via tick bite, we used *Ixodes ricinus* nymphs infected with isolate NE4049 that had been generated in a previous study [30] and that came from our laboratory colony of *Borrelia*-free *I. ricinus* ticks. For these *I. ricinus* nymphs, the percentage of nymphs infected with *B. afzelii* ranged between 80.0% and 100.0%. Uninfected control nymphs were obtained from our laboratory colony of *Borrelia*-free *I. ricinus* ticks.

### Experiment

The experimental design of the study is shown in **Figure 1**. Engorged adult female *I. ricinus* ticks were collected from wild roe deer captured in the Sylve d’Argenson forest near Chizé, France. The female ticks were allowed to lay their eggs in the laboratory. Four weeks after deposition, each of the 10 clutches of eggs was split into two batches. One batch was rinsed with 10% bleach while the other batch was rinsed with water (**Figure 1**). The rinsed eggs were allowed to hatch into larvae under non-sterile conditions. To test whether the bleach treatment had reduced the microbiota in the larval ticks, a group of ∼400 larvae was frozen for each of the 20 batches at six weeks after hatching.

**Figure 1.**
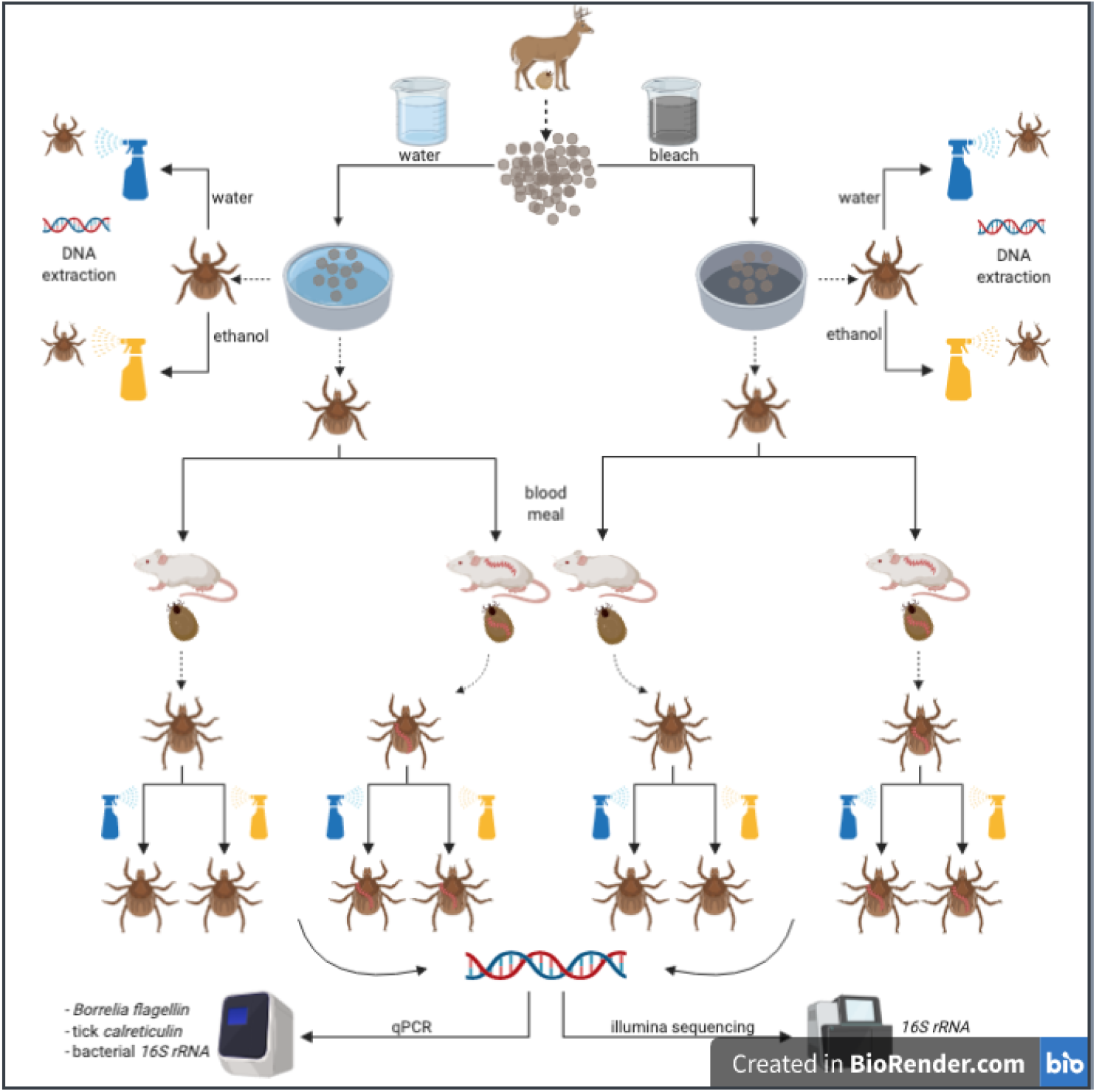
Experimental design. Engorged female *I. ricinus* ticks (n = 10) were collected from roe deer captured in the Chizé forest, France and laid their eggs in the laboratory. Each of the 10 egg batches was split into two batches and rinsed with either 10% bleach (n = 10 batches) or distilled water (n = 10 batches) and hatched into larvae. To determine whether the egg bleaching treatment reduced the microbiome, a subset of larvae was tested using qPCR and Illumina sequencing of the bacterial *16S rRNA* gene. Larvae for each of the 20 batches were split into two groups of ∼100 larvae. For each of the 20 batches of eggs (10 tick families x 2 egg washing treatments), one group of larvae was fed on an uninfected control mouse (n = 20 control mice) whereas the other group of larvae was fed on a *B. afzelii*-infected mouse (n = 20 infected mice). Engorged larvae were placed in individual Eppendorf tubes to moult into nymphs. Four weeks after the moult, 10 nymphs were randomly selected from each of the 40 mice and frozen at – 80°C (n = 400 nymphs). These nymphs were tested for *B. afzelii* infection using qPCR and for their bacterial load and microbiome using qPCR and Illumina sequencing of the bacterial *16S rRNA* gene.

The remaining larvae for each of the 20 batches were split into two groups of ∼100 larvae. For each of the 20 batches of eggs (10 tick families x 2 egg washing treatments), one group of larvae was fed on an uninfected control mouse (n = 20 control mice) whereas the other group of larvae was fed on a *B. afzelii*-infected mouse (n = 20 infected mice; **Figure 1**). These mice had been infected with *B. afzelii* (or not) via nymphal tick bite (see below for details). The resultant engorged larvae were placed in individual Eppendorf tubes and allowed to moult into nymphs. Four weeks after the larva-to-nymph moult, 10 nymphs were randomly selected from each mouse and were frozen at –80°C (**Figure 1**). In summary, we froze 400 nymphs (10 families*2 egg washing treatments* 2 mice infection statuses*10 nymphs/mouse).

### Experimental infection of mice

For the main experiment, 40 BALB/c mice were randomly assigned to either the control group or the infection group. Each mouse in the control group (n = 20) was infested with 5 uninfected *I. ricinus* nymphs, whereas each mouse in the infected group (n = 20) was infested with 5 *B. afzelii*-infected *I. ricinus* nymphs. Each of the 40 mice had been infested with 5 nymphs, so that each mouse had similar immune experience with ticks. Five weeks after the nymphal challenge, an ear tissue biopsy and a blood sample were taken from each of the 40 mice. The ear tissue biopsy was tested for the presence of *B. afzelii* infection using qPCR. The blood sample was tested for *Borrelia*-specific antibodies using a commercial Lyme disease ELISA. These tests confirmed that the 20 mice in the infected group were infected with *B. afzelii*, whereas the 20 mice in the control group were uninfected. These 40 mice were used to feed the larval ticks (see above and in **Figure 1**).

### Molecular methods for larval ticks

The 20 groups of larval ticks that had been frozen were split into two sub-groups with ∼200 larval ticks per sub-group. Half of these sub-groups were washed with ethanol prior to DNA extraction and the other sub-groups were not washed. DNA extraction of the 40 sub-groups of larval ticks was done using a QIAGEN kit following the manufacturer’s instructions. The DNA of each sub-group was eluted into 100 μl of distilled water. The DNA concentration was measured for each of the 40 sub-groups using a Nanodrop. For qPCR, the DNA concentration of each sub-group was adjusted to 5 ng/μl. Two qPCR assays were performed independently for each DNA extraction: tick *calreticulin* and bacterial *16S rRNA*. Each qPCR assay contained 3 μl of template for a total of 15 ng of DNA.

### Molecular methods for nymphal ticks

For each mouse, the 10 nymphs were split into two groups of 5 nymphs. Ticks in one group were washed with ethanol prior to DNA extraction and ticks in the other group were not washed. DNA extraction of whole ticks was done using a QIAGEN kit following the manufacturer’s instructions. The DNA of each tick was eluted into 65 μl of distilled water. The DNA concentration was measured for each of the 400 nymphs using a Nanodrop. For qPCR, the DNA concentration of each tick was adjusted to 5 ng/μl. Three qPCR assays were performed independently for each DNA extraction: tick *calreticulin*, bacterial *16S rRNA*, and *Borrelia flagellin* (see electronic supplementary material (ESM) for details). Each qPCR assay contained 3 μl of template for a total of 15 ng of DNA.

### Illumina Library Preparation and Sequencing of the *16S rRNA* gene

Of the 400 nymphs for which we had quantified bacterial load using the *16S rRNA* gene qPCR assay, Illumina sequencing was performed for 360 nymphs. Sample preparation consisted of two PCR reactions. In the first reaction we amplified a 464 bp fragment of the V3-V4 region of the *16s rRNA* gene using primers Bakt_341F (5’CCTACGGGNGGCWGCAG-3’) and Bakt_805R (5’GACTACNVGGGTATCTAATCC-3’) [33], designed with Illumina adapters. Reactions were performed in a final volume of 50 µl using 2.5 U of HotStar HiFidelity DNA polymerase (Qiagen,Germany), 2.5 µl of 10 µM primers, 10 µl of 15 µM dNTP mix, with a thermal cycle with a denaturation step of 95°C for 5 min, 45 cycles of 94°C for 15 sec, 51°C for 45 seconds, and 72°C for 45 seconds, with a final elongation step at 72°C for 7 minutes. Amplicons were purified with the Wizard SV Gel and PCR Clean-Up system (Promega Switzerland).

The second PCR incorporated the sample barcodes. Reactions were performed in a final volume of 25 µl using 1.25 U of HotStar HiFidelity DNA polymerase, 1 µl of 10 µM primers, 5 µl of 15 µM dNTP mix. The thermocycler had a denaturation step of 95°C for 5 min, 12 cycles of 95°C for 30 sec, 55°C for 30 seconds, and 72°C for 30 seconds, with a final elongation step of 72°C for 5 minutes, and amplicons purified as above. The 360 purified amplicons were pooled in equimolar concentration using a Qubit 2.0 fluorometer (Invitrogen) and sequenced by Microsynth (Balgach, Switzerland) using an Illumina MiSeq v2 with 250 bp paired end output, followed by adaptor and quality trimming.

### Statistical Methods

#### Analysis of the bacterial load in the larvae

For each sub-group of larval ticks, we divided the *16S rRNA* gene copy number by the *calreticulin* gene copy number. These *16S rRNA* to *calreticulin* ratios were log10-transformed to improve the normality of the data. The log10-transformed *16S rRNA* to *calreticulin* ratios were analyzed using linear mixed effects models (LMMs). Fixed factors included egg washing treatment (2 levels: water and bleach), larval tick washing treatment (2 levels: none and ethanol), and their interactions. Random factors included tick family. We used R/Bioconductor (v 3.4.2. or above) for analyses, including the *lme4, complexHeatmap, vegan*, and *phyloseq* packages [34-37].

#### Data selection of the nymphs

Of the 370 DNA extractions, 14 were not included in the analysis because their DNA concentrations were too low. After adjustment of the DNA concentration, the remaining 356 DNA extractions had a DNA concentration that ranged between 3.33 and 5.00 ng/μl so that the 3 μl of DNA template contained between 10.0 and 15.0 ng of DNA.

#### Analysis of the bacterial load in the nymphs

For each tick, we divided the *16S rRNA* gene copy number by the *calreticulin* gene copy number. These ratios were log10-transformed to normalize the data. The log10-transformed *16S rRNA* to *calreticulin* ratios were analysed using LMMs. Fixed factors included egg washing treatment (2 levels: water and bleach), *B. afzelii* infection status of the mouse (2 levels: uninfected control, infected), nymphal tick washing treatment (2 levels: none and ethanol), and their interactions. Random factors included tick family and mouse identity nested inside tick family.

#### Analysis of the *B. afzelii* infection prevalence in the nymphs

These analyses were restricted to the subset of nymphs that had fed as larvae on the *B. afzelii*-infected mice. We used a proportion test to determine whether the microbiome reduction of the egg washing treatment influenced the susceptibility of the nymphs to acquire *B. afzelii* infection during the larval blood meal. The nymphal infection status (0 = uninfected, 1 = infected) was also analysed using generalized linear mixed effects models (GLMMs) with binomial errors. Fixed factors included egg washing treatment, nymphal tick washing treatment, and their interaction. Random factors included tick family and mouse identity nested inside tick family.

#### Analysis of the *B. afzelii* spirochete load in the nymphs

This analysis was restricted to the subset of nymphs that were infected with *B. afzelii*. For each tick, we divided the *B. afzelii flagellin* gene copy number by the tick *calreticulin* gene copy number. These ratios were log10-transformed to normalize the data. The log10-transformed *flagellin* to *calreticulin* ratios were analysed using LMMs. Fixed factors included egg washing treatment, nymphal tick washing treatment, and their interaction. Random factors included tick family and mouse identity nested inside tick family.

#### Analysis of 16S rRNA amplicons

Because of limited overlap between the 250 bp read ends, we elected to use an OTU picking strategy that did not require first assembling paired end reads into contigs. We generated OTU tables using the CD-HIT-OTU-Miseq workflow [38] packaged with CD-HIT v 4.6.8. Forward and reverse read lengths were specified at 200 and 150 bp and clustered against the SILVA 132 99% OTU release [39], otherwise using default parameters that included using Trimmomatic for read trimming [40]. This identified 10,454 OTUs, although the vast majority of reads (93.6%) recruited to the 100 most abundant OTUs. It also assigned taxonomy to only 926 OTUs (< 10%): for more robust taxonomic assignments, we applied Metaxa2 v2.2 [41] to the representative sequences for each OTU identified by CD-HIT using default parameters and the included reference database in Metaxa2. This identified 7,550 OTUs as bacterial and provided taxonomies that were largely congruent with CD-HIT assignments, where evaluable; subsequent analyses were restricted to this bacterial OTU set.

The OTU table and taxonomy were imported to R (>v. 3.4.2), and analyzed using the *phyloseq, vegan*, and *DeSeq2* packages (as detailed in Results). Linear mixed models for univariable outcomes were implemented using the *lme4* package with a family as a random effect, or nesting mouse host within tick family. In analyses using *DeSeq2* and *db-RDA*, family was included as a fixed factor (*DeSeq2*) or a conditioning variable (*db-RDA*). We evaluated the reliability of our replicated sequencing/analysis approach on the same nymphal samples via the variance explained by sample in *db-RDA*, and the intraclass correlation coefficient for Shannon diversity. For statistical analyses requiring tick infection status, we imputed missing infection status for one *Borrelia*-exposed nymph as infected, as that was the most common state of exposed nymphs.

## Results

### Egg bleaching profoundly disrupts the microbiome of tick larvae

Egg bleaching had strong effects on the abundance of bacteria associated with tick larvae, as shown by a 27.5-fold reduction in the relative *16S rRNA* gene (hereafter *16S*) copy number (scored via qPCR) in larvae hatching from bleached versus unbleached eggs (N = 29 evaluable samples; linear mixed model (LMM) controlling for tick family; P < 10^−6^; log_10_ ratio of *16S*/c*alreticulin* 0.24 versus 6.61, respectively; **Figure 2A**). This reduction was evident six weeks after treatment, demonstrating a profound effect of bleaching on the larval tick microbiome. In contrast, washing larvae with ethanol prior to DNA extraction, which is expected to reduce the external microbiota [but see 42], did not have a significant effect on the *16S* copy number (LMM; P = 0.12; **Figure 2A**). Taken together, these observations suggest that the egg bleaching-induced reduction of *16S* copy number was driven by an increase in the relative abundance of the internal microbiota of the larval ticks at the expense of the external microbiota.

**Figure 2.**
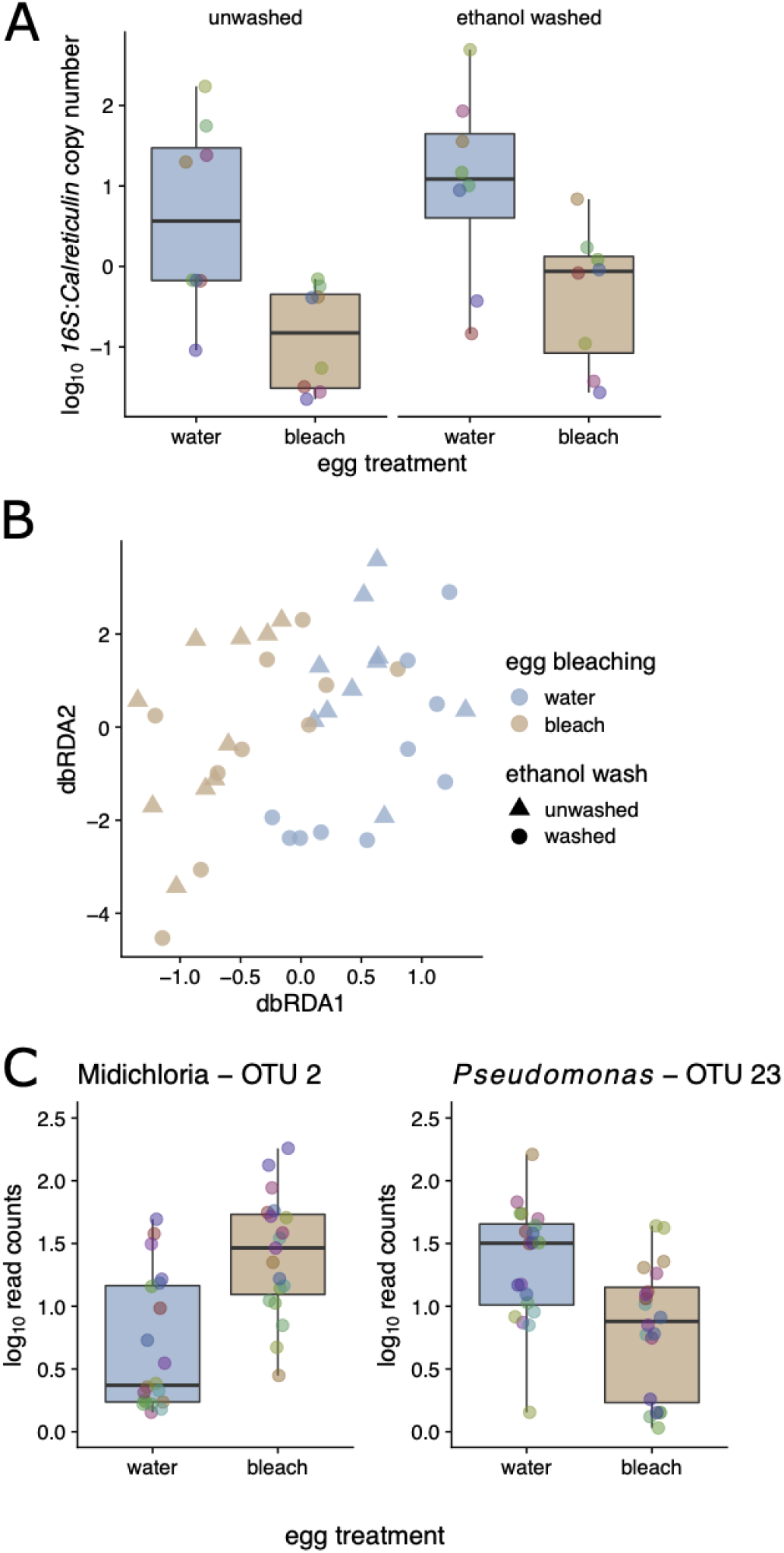
Bleaching of egg casings profoundly decreases the abundance of microbiota in larval ticks. Pooled *Ixodes scapularis* larvae were quantified six weeks after egg bleaching. **A**. Egg bleaching led to a ∼27.5-fold reduction in the relative copy number of the 16S rRNA in larvae (vs tick *calreticulin*; P < 0.001), irrespective of ethanol washing prior to DNA extraction (P > 0.05). **B**. Egg bleaching, but not washing with ethanol prior to DNA extraction, led to significant shifts in the bacterial community, as measured by 16S amplicon sequencing and dbRDA (P < 0.001 and P > 0.05, respectively). **C**. The most enriched taxon in response to egg bleaching was *Candidatus* Midichloria mitochondrii (Padj < 0.001); in contrast the most depleted was a *Pseudomonas* OUT (the Y-axis has units of number of read counts per thousand mapped reads). Colored points in boxplots represent individual data points (pooled larval samples, colored by the tick mother; ‘family’; N = 10).

Multivariate analysis of the larval tick microbiome using db-RDA revealed that egg bleaching led to a clear shift in the *16S* community, whereas there was no discernible effect of washing the larvae prior to DNA extraction (db-RDA; based on Bray-Curtis dissimilarities of log_10_ OTU abundances, stratified by tick family. Permutation tests; P < 0.001 and 0.727, respectively; **Figure 2B**). Consistent with our expectations, the most significantly (proportionally) enriched taxon due to bleaching was in the order Rickettsiales, which we further manually annotated to *Candidatus* Midichloria mitochondrii, an endosymbiont of *I. ricinus* that we expect would be unaffected by external bleaching (**Figure 2C**; P_adj_ < 10^−6^; shown are the log_10_ read counts per thousand mapped reads). In contrast, egg bleaching significantly reduced the relative abundance of Pseudomonas OTU 23 in the resultant larvae, which suggests that this bacterium is found on the surface of the eggshell (**Figure 2C**; P_adj_ < 10^−3^. Other significant effects associated with bleaching included an increase in *Methylobacterium*, a taxon that has been implicated as a potential contaminant of laboratory reagents [43]; these are consistent with the strong reduction in *16S* abundance we observed via qPCR. No OTUs changed significantly from washing (all P_adj_ > 0.05). In sum, egg bleaching dramatically decreased *16S* copy number and shifted the microbial community composition in larvae measured at 6 weeks after treatment. Egg bleaching probably reduced the relative abundance of bacteria associated with the egg surface and thereby increased the relative abundance of endosymbiont bacteria. The lack of an observed effect of washing the larvae—on both microbial abundance and microbial diversity— suggests that external microbiota are a minor component of the *16S* diversity in the lab-hatched tick larvae.

### Manipulation of the larval microbiome has no effect on the acquisition of B. afzelii, and the microbiome largely recovers in the unfed nymphs

We screened nymphs for *B. afzelii* infection via qPCR and found no evidence that disruption of the larval microbiome affected either the percentage of ticks that acquired the infection while feeding (with infection prevalences of 68.2% and 73.5% in unbleached and bleached groups respectively; binomial GLMM; P = 0.44, N = 183, **Figure 3A**) or *B. afzelii* copy number in the nymphs that became infected (LMM; P = 0.958, N = 130; **Figure 3B**). Thus, although the egg bleaching treatment was highly effective at reducing the microbiome in larvae, it did not affect the ability of *B. afzelii* to colonize ticks during or following the larval blood meal.

**Figure 3.**
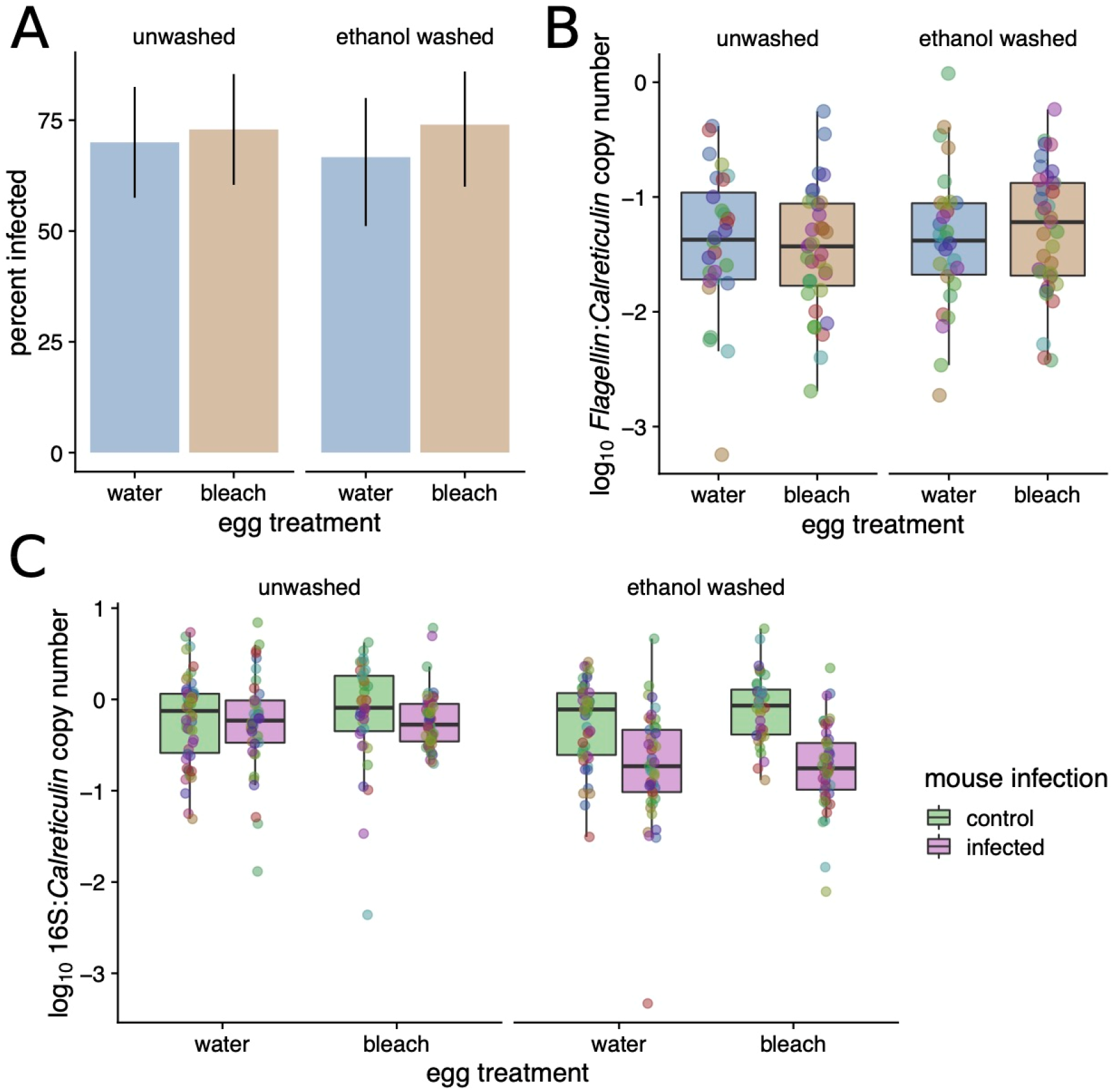
‘Dysbiosing’ larval ticks does not affect infection success by *B. afzelii*. Neither **A**. the percentage of nymphs that acquired *B. afzelii* during their larval blood meal nor **B**. the inferred *B. afzelii* load in these nymphs were affected by prior microbiome disruption (all P >> 0.05). **C**. Feeding on *B. afzelii-*infected mice decreased the bacterial load in *I. ricinus* nymphs; this effect was more visible in the nymphs that were washed prior to DNA extraction compared to the unwashed nymphs.

Although the egg bleaching treatment strongly reduced the *16S* copy number in the larvae, the bacterial community largely recovered after these larvae had taken a blood meal and molted into nymphs, as there was no effect of the egg bleaching on *16S* copy number in nymphs (LMM; P = 0.272, N = 356, **Figure 3C**). Intriguingly, in contrast to the non-significant effect of ethanol washing on *16S* copy number that we observed in larvae, there was a clear reduction with both washing and *B. afzelii* exposure. The *16S* copy number was lowest in ticks that were both washed and *B. afzelii*-exposed (LRT for interaction; P < 10^−5^), indicating that feeding on *B. afzelii-*infected mice reduced the internal bacterial load in nymphs; we investigate this hypothesis in more detail below.

### Host infection with B. afzelii has pervasive effects on the microbiome of I. ricinus nymphs

To extend our analysis of larval ticks to the nymphal stage, we used *16S* amplicon sequencing to profile replicate nymphs from each treatment alongside the larval samples (above). We again restricted these analyses to the 40 most abundant OTUs. As expected from prior reports [44, 45], there was a clear shift in the tick-associated bacterial community from larval to nymphal ticks (**Figure 4A**). There was, further, a striking effect of *B. afzelii* infection in the mouse on the tick microbiota as quantified in the nymphal stage, with pronounced differences between ticks fed on *B. afzelii*-infected versus control mice (db-RDA; P < 0.001; **Figure 4B** shows unsupervised principal coordinates analysis (PCoA)). We also found that egg bleaching and ethanol washing prior to DNA extraction had modest but significant effects on microbiome composition in nymphs (P < 0.001 and P = 0.014, respectively; **Figure 4B**), consistent with our finding that washing the nymphs decreased *16S* copy number. While sequencing recovered OTUs annotated as *Borrelia*, the most abundant *Borrelia-*annotated OTU was ranked 81^st^ in overall abundance and accounted for <0.05% of total sequence reads. Thus, a direct contribution of sequenced *Borrelia 16S* amplicons in infected ticks does not explain these strong patterns.

**Figure 4.**
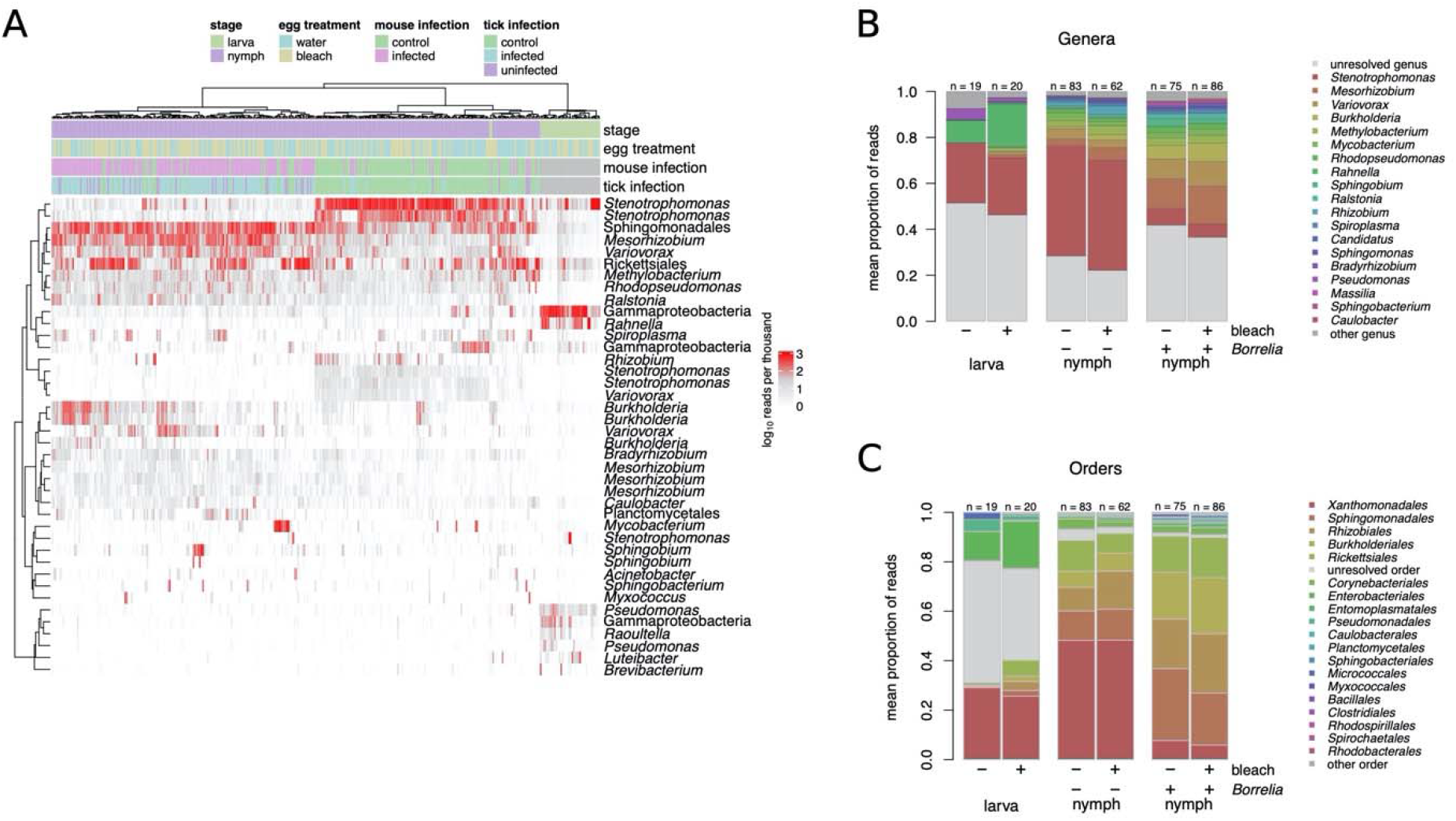
Life history stage and *B. afzelii* exposure affect tick microbiome. **A**. Heatmap of number of reads assigned (log_10_(x+1) per thousand) for top 40 OTUs across all samples in the experiment. Highest taxonomy reliably assigned by Metaxa2 is shown. Dendrograms are based on hierarchical clustering of Bray-Curtis dissimilarities using Ward’s method. **B**. Composition of treatment groups and life histories, with top 40 OTUs aggregated (as mean of samples per group) at the genus level. **C**. As above, at the order level.

Microbiome α-diversity (measured as Shannon entropy) in nymphs modestly increased as a result of both microbiome disruption via egg bleaching (LMM; P = 0.035; N = 305. **Figure 5A**) and from *B. afzelii* infection (P = 0.0025; **Figure 5B**) or exposure (P = 0.0007; competing multivariable model; **Figure 5B**). Coupled with the decrease in *16S* copy number observed when feeding on *B. afzellii-*infected mice, these diversity effects appeared to be mediated through disproportionately negative impacts on abundant OTUs (e.g., *Stenotrophomonas*) leading to increased community evenness. As mentioned, these trends were evident when considering the infection status of the tick itself, or that of the mouse on which they were fed (**Figure 5C**). Comparison between control nymphs, uninfected nymphs, and infected nymphs demonstrated that it was feeding on an infected mouse rather than acquiring *B. afzelii* infection that was most important for determining the nymphal microbiome (**Figure 5C**). This was supported by comparing competing models with Akaike’s Information Criterion (AIC), which found that feeding on an infected mouse was a much stronger predictor of both α-diversity (ΔAIC ∼ 6) and the multivariate bacterial community (**Figure 5A;** ΔAIC ∼14 in models using first PCoA axis as response variable) than acquisition of *B. afzelii* by the tick (**Figure 5**), suggesting that the effects we observe are more likely to be caused by physiological or immunological characteristics of infected mice rather than the direct effects of *B. afzelii* infection in the ticks. Similarly, examining the OTU frequencies in bleached and *B. afzelii*-exposed ticks showed that these net effects were driven by proportional reductions in the dominant OTUs, with concomitant increases in less frequent OTUs (e.g. **Figure 4**). Collectively, these results are consistent with the strong effects of mouse *B. afzelii* infection status we observe on *16S* copy number and suggest that these are specifically mediated by disproportionate negative effects on abundant bacterial OTUs.

**Figure 5.**
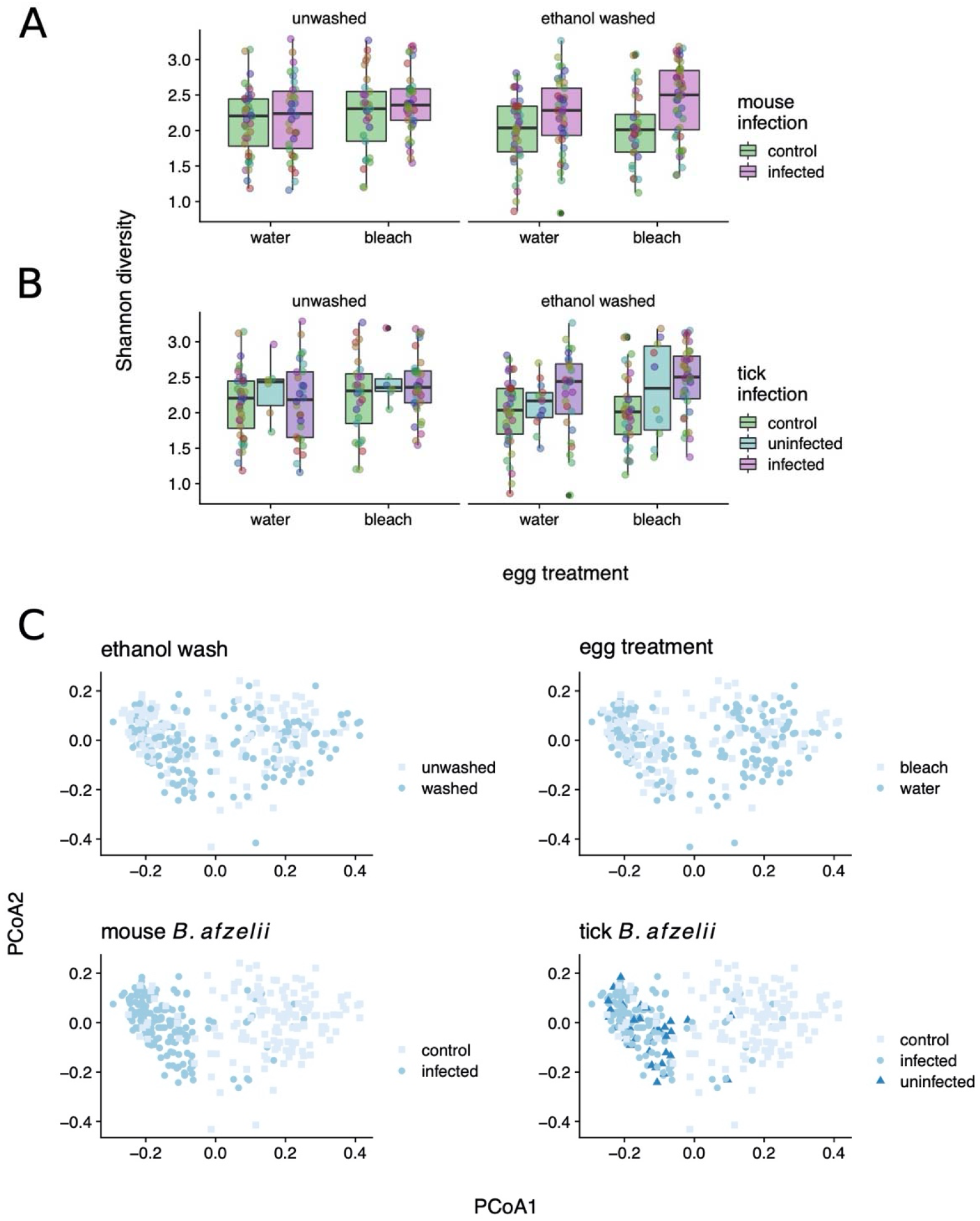
Egg bleaching and rinsing and *B. afzelii* exposure increase the diversity of the 16S microbiome of *I ricinus* nymphs. Shannon diversity increases in both **A**. *B. afzelii* exposed ticks and **B**. *B. afzellii* infected ticks (P <0.01). Akaike’s Information Criterion (AIC) shows exposure to be a stronger predictor of Shannon diversity than infection (deltaAIC ∼6) in linear mixed models. Points represent individual tick nymphs colored by family of origin. C. Principal Coordinates Analysis of nymphal 16S counts, colored by experimental factors; *B afzelii* exposure best stratifies groups on PCoA 1 (deltaAIC ∼11 vs infection).

### Microbial correlates of Borrelia exposure

To characterize the recurrent shifts in the tick microbiome associated with *B. afzelii* exposure, we used negative binomial models implemented in *phyloseq/DESeq2*. In line with the global diversity shifts we observed, feeding on *B. afzelii-*infected mice led to significant changes in the relative frequencies of many OTUs (with 19/40 significant at P_adj_ < 0.05; Figure 5). This was most evident in large decreases in the frequency of multiple *Stenotrophomonas* OTUs and other Gammaproteobacteria (**Figure 6**). As a result, there appeared to be a degree of taxonomic dependence in microbial responses to *B. afzelii* exposure, with Betaproteobacteria generally significantly increasing in frequency and Gammaproteobacteria decreasing (Fisher’s exact test, P = 0.004; **Figure 6)**. Our analysis also revealed that several nymph OTUs that increased in frequency in response to tick bleaching, including two *Burkholderia* OTUs (Betaproteobacteria; P_adj_ < 0.01; **Figure 6**) and a *Bradyrhizobium* OTU (Alphaproteobacteria; P_adj_ < 0.01; **Figure 6**), whereas no OTUs significantly (proportionally) decreased. Consistent with our expectations, ethanol washing prior to DNA extraction increased the relative abundance of the endosymbiotic *Candidatus* Midichloria mitochondrii (annotated as order *Rickettsiales*, Alphaproteobacteria) as well as *Spiroplasma* (Mollicutes) by 2.5 and 1.7-fold respectively (P_adj_ < 0.01), corroborating an enriching effect of washing on the internal tick microbiota.

**Figure 6.**
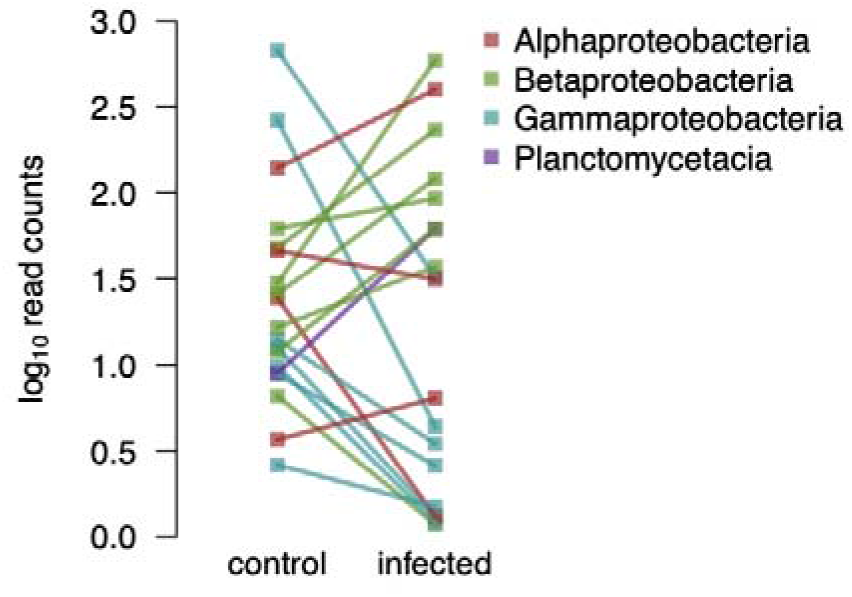
Proportional abundance of taxa that changed significantly in *I. ricinus* nymphs (19/40 at P_adj_ < 0.05) in response to feeding on *B. afzelii* infected mice, shown at the level of class. There is significant taxonomic dependence (Fisher’s exact test P = 0.004) of taxonomic response.

## Discussion

Here, we used a highly replicated experiment on wild-collected tick families to examine the reciprocal interactions between the endogenous tick microbiota and *B*.*afzelii*. We used egg bleaching to radically disrupt the microbiome in larval ticks but found no evidence that this affected the subsequent susceptibility of ticks to infection with *B. afzelii*. Rather, this work revealed striking effects of feeding on *B. afzelii-*infected mice on the tick microbiome; these effects superseded those of actually acquiring an infection, as judged by comparing competing statistical models, and the resultant partitioning of samples in microbial community space (e.g., **Figure 4A, 5C**).

A growing number of studies have investigated whether dysbiosing ticks influences their susceptibility to acquiring tick-borne pathogens [8, 11-13]. In some systems, microbiome disruption makes tick species more susceptible to infection with tick-borne pathogens [8, 13], whereas other systems found the opposite effect [11, 12]. In the present study, we found no evidence that microbiome disruption influenced *B. afzelii* infection rates or the *B. afzelii* spirochete load in infected ticks. In contrast, a previous study found that dysbiosed *I. scapularis* larvae were less likely to acquire *B. burgdorferi* ss and contained lower bacterial loads compared to control larvae [12]. These two studies differed in a number of factors including the *Borrelia* species, the *Ixodes* tick species, and the method of dysbiosis. An important aspect of the present study is that we investigated whether dysbiosis of the eggs influenced the infection status of the nymphs, which is the stage that is actually critical for the transmission of Lyme disease in nature [46].

Despite the fact that bleaching the eggs was highly effective at reducing the bacterial microbiome in the resultant larvae, this method of ‘dysbiosis’ did not have a meaningful impact on *B. afzelii* transmission. While the presence of additional bacteria to those uncovered here could influence these patterns, we found little influence of tick family on the recovered bacterial communities, outside of potentially vertically transmitted bacteria such as *Spiroplasma*, suggesting the microbial community of *I. ricinus* is largely homogenous at the scale studied here. Our method of dysbiosis further highlights the neglected importance of maternal transmission of gut symbionts in ticks. In arthropods, gut symbionts are typically vertically transmitted by superficial bacterial contamination of eggs (egg smearing) [47]. Our egg washing with bleach has removed such maternally inherited gut symbionts and this impacted the microbial communities hosted by larvae. Our study shows that egg smearing is a key mechanism for colonization of ticks by their associated microbes.

*B. afzelii* infection reduced the microbial abundance (in the ethanol-washed nymphs) and changed the microbial community in the unfed nymphs. Notably, we observed changes in OTU relative abundance to be, at least in part, taxon-specific with decreases in Gammaproteobacteria and increases in Betaproteobacteria (**Figure 6**). The observation that mouse infection status was more important than tick infection status suggests that the blood physiology at the time of the larval blood meal was critical for structuring the subsequent nymph microbiome. Metabolomic studies of mouse serum samples have shown that *B. burgdorferi* ss infection changes the blood concentration of amino acids, energy metabolites, and aromatic compounds [48], which could influence the development of the tick microbiome. Infection with *B. burgdorferi* sl stimulates the host immune system, which could also exert collateral damage on the tick microbiome [49-52]. For example, elevated levels of complement, cytokines, leukocytes, and reactive oxygen species in the blood [52-55] may interact inside the tick to have negative effects on the midgut microbiome. In summary, our study suggests that the physiological and immunological changes associated with infection in the vertebrate host have important consequences for the microbiome of feeding ticks.

## Conclusions

In summary, we found that egg bleaching resulted in a 30-fold reduction of the microbiome of larval ticks. This microbiome manipulation had no effect on the ability of larval ticks to acquire *B. afzelii* after feeding on infected mice. Once the engorged larvae had moulted into unfed nymphs, the dramatic effect of the egg bleach treatment on the tick microbiome had mostly disappeared. The *B. afzelii* infection status of the mice that provided the larval blood meal had a dramatic effect on the microbiome of the resultant unfed nymphs. Our study suggests that infection in the vertebrate host influences the quality of the larval blood meal with long-term consequences for the tick microbiome that persist into the nymphal stage.

## Ethics approval and consent to participate

The commission that is part of the “Service de la Consommation et des Affaires Vétérinaires (SCAV)” of Canton Vaud, Switzerland evaluated and approved the ethics of this study. The Veterinary Service of the Canton of Neuchâtel, Switzerland issued the animal experimentation permit used in this study (NE04/2014).

## Availability of data and materials

Raw sequencing reads will be deposited at NCBI (accession pending). Supplementary data and scripts to reproduce the analysis will be made available at github.com/onecarbon/tickdysbiosis (pending).

## Competing interests

The authors declare that they have no competing interests.

## Funding

This work was supported by the following grants awarded to Maarten J. Voordouw: a Swiss National Science Foundation grant (FN 31003A_141153) and a Discovery Grant from the Natural Sciences and Engineering Research Council of Canada (RGPIN-2019-04483). PTH was supported by a Canadian Institutes for Health Research PDF.

## Authors’ contributions

EM, OD, OP and MJV conceived and designed the study. EM, AS, and AB conducted the experiment and performed the molecular work. PTH and MJV analysed the data and wrote the manuscript. GH created the figure of the experimental design. All authors read and approved the final version of the manuscript.

## Acknowledgments

We thank Gilles Capron (Office National de la Chasse et de la Faune Sauvage) and the Office National des Forêts of the Réserve biologique domaniale intégrale de la Sylve d’Argenson for permission to collect ticks on roe deer captured in the Chizé forest (France).

## Supplemental Material

**Data S0:** Description of molecular methods

**Data S1:** Metadata

**Data S2:** Metadata

**Data S3:** OTU table produced by CD-HIT-OTU_MiSEQ. **Data S4:** OTU clusters produced by CD-HIT-OTU_MiSEQ. **Data S5:** OTI taxonomy assigned by Metaxa2.

## Notes

### Competing Interest Statement

The authors have declared no competing interest.

