## Supplementary Methods for "*Borrelia* infection in rodent host has dramatic effects on the microbiome of ticks"

***Borrelia flagellin* gene qPCR:** The total spirochete load of *Borrelia afzelii* in the *Ixodes ricinus* nymphs was estimated using a qPCR that targeted a 132-bp fragment of the *flagellin* gene (Schwaiger et al 2001). The qPCRs were performed using the LightCycler^®^ 480 Multiwell Plate 96 white (Roche). The wells were filled with a mixture of 5.8 µl of water, 10.0 µl of Master Mix (FastStart Essential DNA probes Master, Roche), 0.4 µl of 20.0 µM forward primer FlaF1A, 0.4 µl of 20.0 µM reverse primer FlaR1, 0.4 µl of 10.0 µM Flaprobe1, and 3.0 µl of DNA template. The thermocycling conditions consisted of 10 min at 95°C for denaturation, followed by 50 cycles of 30 sec at 60°C and 10 sec at 95°C.

**Bacterial *16S rRNA* gene qPCR**: We quantified bacterial load in each tick using a SYBR Green real-time qPCR assay that amplified the V3 hypervariable region of the 16S rRNA gene. The primers used were 338f (5’-ACTCCTACGGGAGGCAGCAG-3’) and 520r (5’-ATTACCGCGGCTGCTGG-3’) (Bakke et al 2011, Muyzer et al 1993). The qPCR was based on a previously developed protocol (Bueche et al 2013). The qPCRs were carried out in a final reaction volume of 10 µl with 5 µl Rotor-Gene SYBR green PCR master mix (Qiagen GmbH, Hilden, Germany), 0.30 µM of forward primer 338f, 0.30 µM of reverse primer 520r, and and 3.0 µl of DNA template. The thermocycling conditions consisted of an initial denaturation/activation step at 95°C for 15 min, followed by 40 cycles composed of denaturation at 95°C for 10 s, annealing at 55°C for 15 s, and elongation at 72°C for 20 s. Each 96-well qPCR plate contained 5 standards, and 3 negative controls that were all run in triplicate. The five standards contained 10^7^, 10^6^, 10^5^, 10^4^ and 10^3^ copies of the *16s rRNA* gene.

Standards were produced using a plasmid containing the *16s rRNA* gene sequence (Bueche et al 2013). Plasmid-transformed *E. coli* cells were grown up overnight and plasmid DNA was extracted with the Wizard Plus SV Miniprep DNA purification system (Promega, Switzerland) following the manufacturer’s instructions. The DNA concentration of the plasmid mini-prep was estimated using a Nanodrop 2000 (Thermo Scientific) and the number of 16s rRNA gene copies was calculated using the known molecular weight of the plasmid (3326251.8 g/mol).

***Ixodes ricinus calreticulin* gene qPCR:** Variation in DNA extraction efficiency and DNA concentration in the DNA extractions will influence the estimates of the *flagellin* gene copy number and the *16S rRNA* gene copy number in the samples. To control for this variation, we standardized our estimates of bacterial abundance with respect to an estimate of the *I. ricinus* nuclear *calreticulin* gene copy number in the samples. We used a SYBR Green real-time qPCR assay that targeted a 109-bp fragment of the *calreticulin* (*cal*)gene as previously described (Sassera et al 2008). The primers used were calF (5’- ATCTCCAATTTCGGTCCGGT -3’) and calR (5’- TGAAAGTTCCCTGCTCGCTT -3’) (Sassera et al 2008). The qPCRs were carried out in a final reaction volume of 20 µl with 12.5 µl Rotor-Gene SYBR green PCR master mix (Qiagen GmbH, Hilden, Germany), 0.40 µM of forward primer calF, 0.40 µM of reverse primer calR, and and 3.0 µl of DNA template. PCR cycling conditions for the *calreticulin* gene were as follows: 95°C for 2 min, 40 cycles at 95°C for 15 s and at 60°C for 30 s, and melt curve analysis from 55°C to 95°C with increasing increments of 0.5°C per cycle.

Each 96-well qPCR plate contained 5 standards, and 3 negative controls that were all run in triplicate. The five standards contained 10^7^, 10^6^, 10^5^, 10^4^ and 10^3^ copies of the *calreticulin* gene. Standards were produced using a pGEMT-easy plasmid containing the *calreticulin* gene sequence. Plasmid-transformed *E. coli* cells were grown up overnight and plasmid DNA was extracted with the Wizard Plus SV Miniprep DNA purification system (Promega, Switzerland) following the manufacturer’s instructions. The DNA concentration of the plasmid mini-prep was estimated using a Nanodrop 2000 (Thermo Scientific) and the number of *calreticulin* gene copies was calculated using the known molecular weight of the plasmid.

**Repeatability of the 16S microbiome:** To test the reproducibility of our estimates of the tick microbiome, we performed replicate library preparation and *16S* sequencing on the 40 DNA extractions of the larval ticks (10 families ✕ 2 egg washing treatments ✕ 2 larva washing treatments prior to DNA extraction), and clustered sequences into operational taxonomic units (OTUs) at 97% sequence identity. After quality control with >500 reads passing QC, 33 of the 40 samples were replicated. We found that metrics derived from duplicate samples were largely reproducible; for example Shannon diversity (see below) showed an intraclass correlation coefficient of 0.82 across replicates, and distance-based redundancy analysis (db-RDA; based on Bray-Curtis Dissimilarity) revealed that 85% of the variance in multivariate community structure was attributable to sample. For subsequent analyses, we summed replicate OTU counts for each larval sample where available, and (with the exception of diversity metrics) restricted our analysis to the top 40 most-abundant OTUs present across our entire experiment; these encompassed ~90% of sequenced reads (including OTUs recovered in nymphs; see below. Cumulative reads per OTU shown in **Figure S1**).

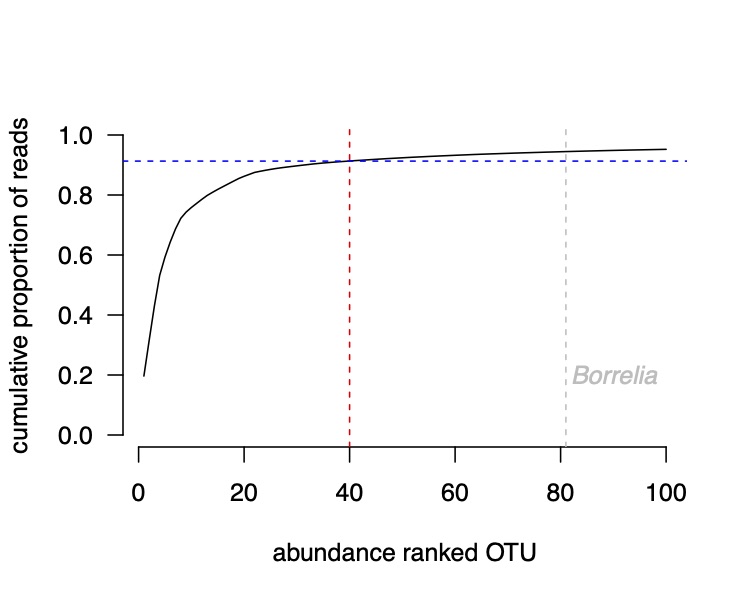

**Figure S1.** Cumulative abundance of abundance-ranked OTUs assayed. The top 40 OTUs comprise >90% of recovered sequences, and the most abundant *Borrelia* OTU identified is 81st in abundance comprising <0.05% of total reads.
